# The silencing factor Sir3 is a molecular bridge that sticks together distant loci

**DOI:** 10.1101/2020.06.29.178368

**Authors:** Myriam Ruault, Vittore F. Scolari, Luciana Lazar-Stefanita, Antoine Hocher, Isabelle Loïodice, Camille Noûs, Romain Koszul, Angela Taddei

## Abstract

Physical contacts between distant loci contribute to regulate genome function. However, the molecular mechanisms responsible for settling and maintaining such interactions remain poorly understood. Here we investigate the well conserved interactions between heterochromatin loci. In budding yeast, the 32 telomeres cluster in 3-5 foci in exponentially growing cells. This clustering is functionally linked to the formation of heterochromatin in subtelomeric regions through the recruitment of the silencing complex SIR composed of Sir2/3/4. Combining microscopy and Hi-C on strains expressing different alleles of *SIR3*, we show that the binding of Sir3 directly promotes long range contacts between distant regions, including the rDNA, telomeres, and internal Sir3 bound sites. Furthermore, we unveil a new property of Sir3 in promoting rDNA compaction. Finally, using a synthetic approach we demonstrate that Sir3 can bond loci belonging to different chromosomes together, when targeted to these loci, independently of its interaction with its known partners (Rap1, and Sir4), Sir2 activity or chromosome context. Altogether these data suggest that Sir3 represents an uncommon example of protein able to bridge directly distant loci.

## Introduction

The physical organization of the genome in space and time can potentially impact its functions. However, the mechanisms underlying this organization and its dynamics remain poorly understood. Among the physical principles governing the 3D folding of genome, contacts between distant loci represent a high potential to regulate biological functions as exemplified by the enhancer-promoter interactions in metazoan. Another example observed from yeast to man is provided by the clustering of repeated sequences sequestering silencing factors and thus forming repressive subcompartments (Meister and Taddei 2013).

The formation of those subcompartments in the nuclear space has several functional consequences. First, it allows a more robust and specific repression of the associated sequences. Subcompartments allow a local concentration of silencing factors, and consecutively, sequester these factors from the outside, limiting their action elsewhere in the genome. Second, the spatial proximity of the clustered sequences favor recombination events between theses sequences (Batté et al. 2017). Finally, since these sequences can be located on different chromosomes or at long distance on the same chromosome, subcompartments affect the global genome folding in 3D space. Therefore, understanding the molecular mechanisms driving and regulating these long-range interactions provides insights on several DNA mechanisms, including genetic regulation, the maintenance of chromosomal stability, or also compaction and folding throughout the cell cycle. Chromatin tethering or co-localization involve a variety of direct or indirect molecular mechanisms, such as cohesin-mediated loop formation (Nasmyth 2005; Rao et al. 2014), or anchoring of heterochromatin to the nuclear lamina (Falk et al. 2019). However, protein-mediated, direct bridging of distant loci positioned on the same or different chromosomes remain scantily described experimentally.

In exponentially growing cells, the 32 telomeres of *S. cerevisiae* cluster within 3 to 5 foci mainly found at the nuclear periphery, where the silencing information regulators Sir2, Sir3 and Sir4, forming the SIR complex, concentrate (Gotta et al. 1996). These clusters represent a well-documented example of repressive nuclear subcompartment (Gartenberg and Smith 2016; Kueng et al. 2013). This organization is regulated by the physiological state of the cells, as illustrated in long lived quiescent cells, in which telomeres cluster into a unique, large focus in the center of the nucleus (Guidi et al. 2015).

At the molecular level, the SIR complex is recruited at the telomeric and sub-telomeric TG1-3 repeats by the binding of the transcription factor Rap1, whose C-terminal end contains binding domains for the silencing factors Sir3 and Sir4 (Gartenberg and Smith 2016). The SIR complex is also recruited at the silent mating type loci (*HM* loci), through different DNA binding proteins, which like Rap1 have other functions in the cell such as Orc1 and Abf1 (Haber 2012). In addition, Sir2 is found at the rDNA locus where it protects rDNA repeats from recombination. Although Sir3 was initially thought to associate to the nucleolus only in aged cells (Kennedy et al. 1997; Sinclair 1997), it was later found to interact with the rDNA by chromatin immunoprecipitation (ChIP) in exponentially growing cells (Radman-Livaja et al. 2011). Furthermore, genome-wide ChIP experiments revealed that Sir3 can also be found at a handful of discrete, telomere-distal, sites (Hocher et al. 2018; Mitsumori et al. 2016; Radman-Livaja et al. 2011; Sperling and Grunstein 2009; Takahashi et al. 2011; Teytelman et al. 2013). Once recruited the SIR complex has the ability to spread along the chromatin fiber and to repress the transcription of underlying genes by the RNA polymerase II. This spreading results from the association of the histone deacetylase activity of Sir2 with the affinity of Sir3 for unmodified nucleosomes, linked by Sir4 interactions with both Sir2 and Sir3. However, SIR spreading is limited by other histone modifications including H3K79me3, which prevents Sir3 binding in euchromatic regions and by the titration of SIR proteins, especially of Sir3. Indeed, overexpression of Sir3 leads to the formation of extended silent domains (ESD) restricted to subtelomeric regions and delimited by transition zones where H3K79me3 levels increases abruptly (Hocher et al. 2018).

A consequence of the limiting amount of SIR proteins is that loci associated with these proteins compete for these limiting pools (Buck and Shore 1995; Michel 2005; Smith et al. 1998). Telomere clustering by increasing the local concentration within these subcompartments could thus favor SIR spreading within subtelomeric regions. Reciprocally, telomere clustering depends on SIR recruitment at telomeres (Gotta et al. 1996). Furthermore, Sir3 overexpression leads to increased telomere clustering in addition to increased SIR spreading (Ruault et al. 2011). A Sir3 point mutation (A2Q) enabled to disentangle telomere clustering and SIR spreading. This mutant has lost its ability to bind nucleosomes and thus to spread along subtelomeric regions but yet can promote telomere clustering (Hocher et al. 2018; Ruault et al. 2011; Sampath et al. 2009). This led us to propose that Sir3 is the bridging factor between telomeres (Meister and Taddei 2013; Ruault et al. 2011).

Here we directly test this hypothesis by exploring the role of Sir3 on the global organization of the genome by combining genome-wide capture of chromosome conformation (Hi-C) and microscopy. We show that Sir3 bound regions tend to contact each other independently of their chromosome location, as we observe preferential interactions between the rDNA, telomeres, and internal Sir3 bound sites. Finally, we show that tethering of Sir3 alone at two ectopic sites belonging to different chromosomes is sufficient to bridge these loci, showing that Sir3 acts as a molecular “glue” to bridge chromatin loci together.

## RESULTS

### Sir3 impacts on genome organization

To explore the influence of Sir3 on the 3D organization of the genome we performed Hi-C experiments in the absence or presence of different *sir3* alleles. We previously showed that Sir3 is a limiting factor for telomeres clustering. Indeed, while the telomere-bound protein Rap1 fused to GFP in living cells forms 3-5 foci at the nuclear periphery in wild-type cells, only weak residual foci can be detected in *sir3*Δ cells possibly resulting from the random encounter of telomeres (Figure 1A and (Ruault et al. 2011)). On the contrary, *SIR3* overexpression leads to the grouping of telomeres in larger foci, or hyperclusters, located in the nuclear interior (Ruault et al. 2011). Moreover, overexpression of the spreading deficient *sir3-A2Q* mutant, which contains a N-terminal substitution blocking its acetylation and thus silencing (Wang et al. 2004) and spreading in subtelomeric regions (Hocher et al. 2018), also resulted in the formation of telomere hyperclusters ((Ruault et al. 2011) and Figure 1A). In addition, whereas telomere clusters tend to dissociate in wild-type during mitosis (Figure 1A), *SIR3* overexpression mediated-hyperclusters persist during this phase of the cell cycle (right panel of Figure 1A). The drastic spatial reorganization of telomeres may influence, directly or not, the global genome organization. To tackle this question, we applied Hi-C to probe the average 3D organization of the genome of the above strains (Lieberman-Aiden et al. 2009). To prevent unwanted influence of the loop organization of mitotic chromosomes characterized in cycling cells (Dauban et al. 2020; Garcia-Luis et al. 2019), Hi-C libraries were obtained from G1, elutriated daughter cells (Methods). The contact maps of wild-type cells revealed an enrichment of contacts between telomeres (red dots on the map) as well as between centromeres as reported previously (Duan et al. 2010; Guidi et al. 2015; Lazar-Stefanita et al. 2017). The enrichment in telomere-telomere contacts disappears in the absence of Sir3 (*sir3*Δ background; Figure 1B, black arrowheads, see Figure S1 for higher resolution maps), whereas centromere-centromere contacts remain apparent (black * on the map). In the strain overexpressing *SIR3*, inter-telomere contact frequency was strongly increased (Figure 1B) to the detriment of contacts with other part of the genome (white lines in between red dots), while frequency of inter-centromere contacts appeared unchanged compared to a WT strain (black * on the map). These differences can be visualized more quantitatively by computing the log2-ratio plot between mutants and WT maps (Figure 1C). In these maps, the color scale reflects the balance of contacts between the two conditions: the bluer, the more contacts in the WT; the redder, the more contacts in the mutant. The comparison of WT and *sir3*Δ maps results in the visualization of inter-subtelomeres contact ratio as blue pixels, as expected from the drop in inter-subtelomere contact frequency observed in *sir3*Δ strain compared to WT. On the opposite, the red stripes expanding from, and bridging subtelomeric regions indicate that the declustered telomeres are now able to “see” more other regions of the genome in the absence of Sir3, reflecting that telomeres are less constrained in the absence of clustering (see also (Muller et al. 2018) for a similar effect during meiosis prophase).

**Figure 1:**
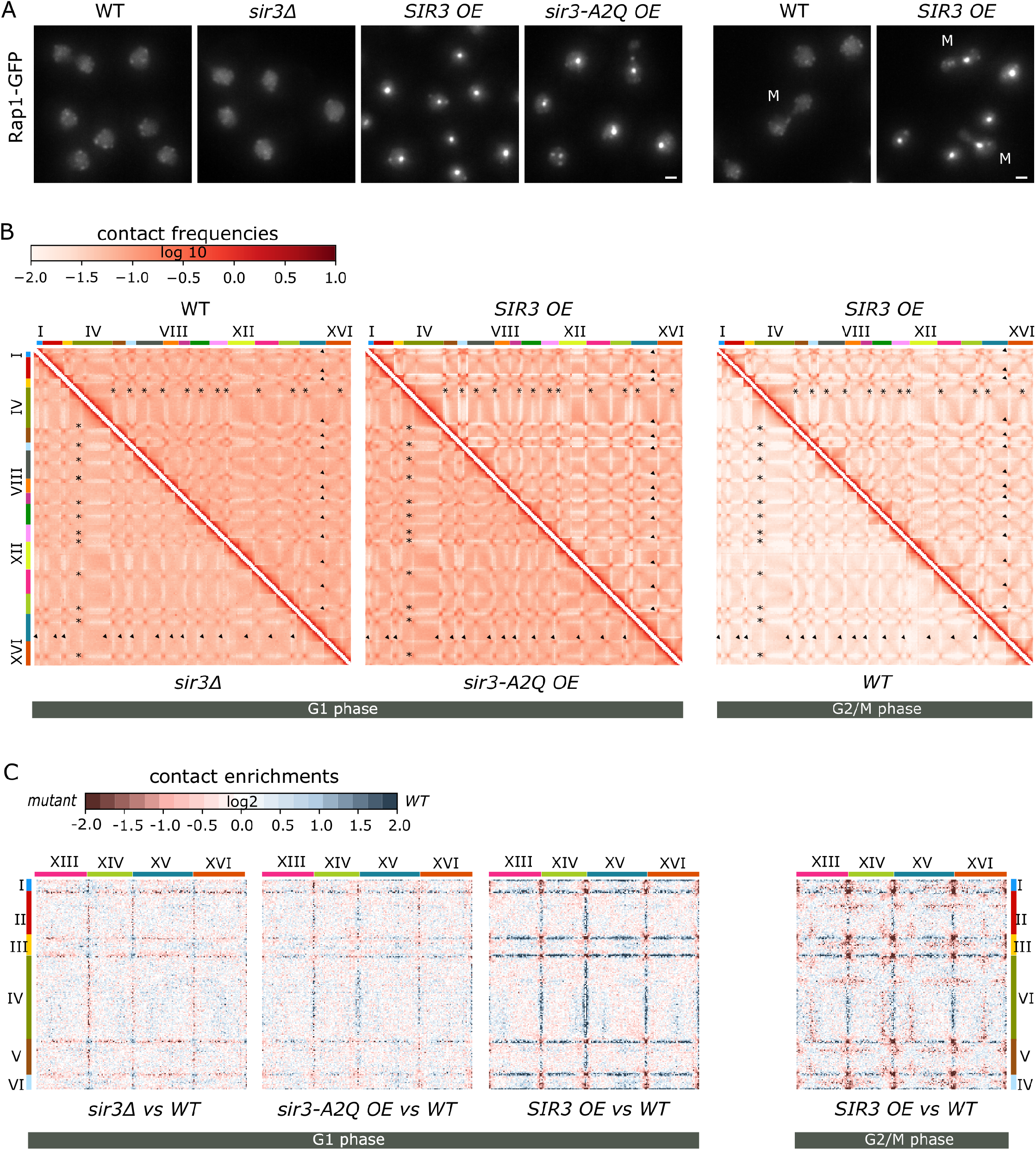
Sir3 impact in genome organization. **(A)** Representative fluorescent images of the telomere-associated protein Rap1 tagged with GFP in exponentially growing strains expressing different Sir3 levels: wild-type (yAT2583), high Sir3 levels (yAT2476), high Sir3-A2Q levels (yAT2822) or no Sir3 (yAT2584). Right two panels: representative mitotic cells (M) in the wild-type strain or in the strain overexpressing Sir3. (**B**) Normalized chromosome contact maps (bin: 50 kb) of cells expressing various levels of Sir3 synchronized either in G1 or G2/M. (**C**) Ratio plots of pairs of contact maps of four representative chromosomes. Blue: enrichment of contacts in wt. Red: enrichment in mutants.

In strains overexpressing the silencing dead Sir3-A2Q protein, the frequency of inter-telomere contacts also increased compared to WT, in agreement with the microscopy, although not to the extent of the strains overexpressing Sir3. The ratio map of the Sir3-A2Q overexpressing cells versus WT cells show an inverted pattern compared to the *sir3*Δ vs WT, with red dots at the level of the telomeres and blue lines in between telomeres (Figure 1C). A similar but stronger pattern was observed when comparing WT and Sir3 overexpressing strains in G1. This pattern was even more contrasted for G2/M cells, as telomere contacts are barely detectable in the wild-type strain while they remained high upon Sir3 overexpression (Figure 1B and 1C right panels). Altogether, these data showed that contacts between subtelomeres decrease in the absence of Sir3, while they increased upon Sir3 or Sir3-A2Q overexpression, in good agreement with the microscopy data.

### Sir3 binding coincides with inter-subtelomere contacts

To determine whether Sir3 overexpression has an impact on global chromosome compaction, we plotted the contact probability curve, *p*(*s*). As expected, the curves display an enrichment in contacts for distances ~10 kb to 100 kb in G2/M compared to G1 samples, reflecting the condensation of chromosomes during mitosis (Lazar-Stefanita et al. 2017). However, different amount of Sir3 did not modify significantly the compaction of the genome measured at the global level in G1 phase (Figure 2A).

**Figure 2:**
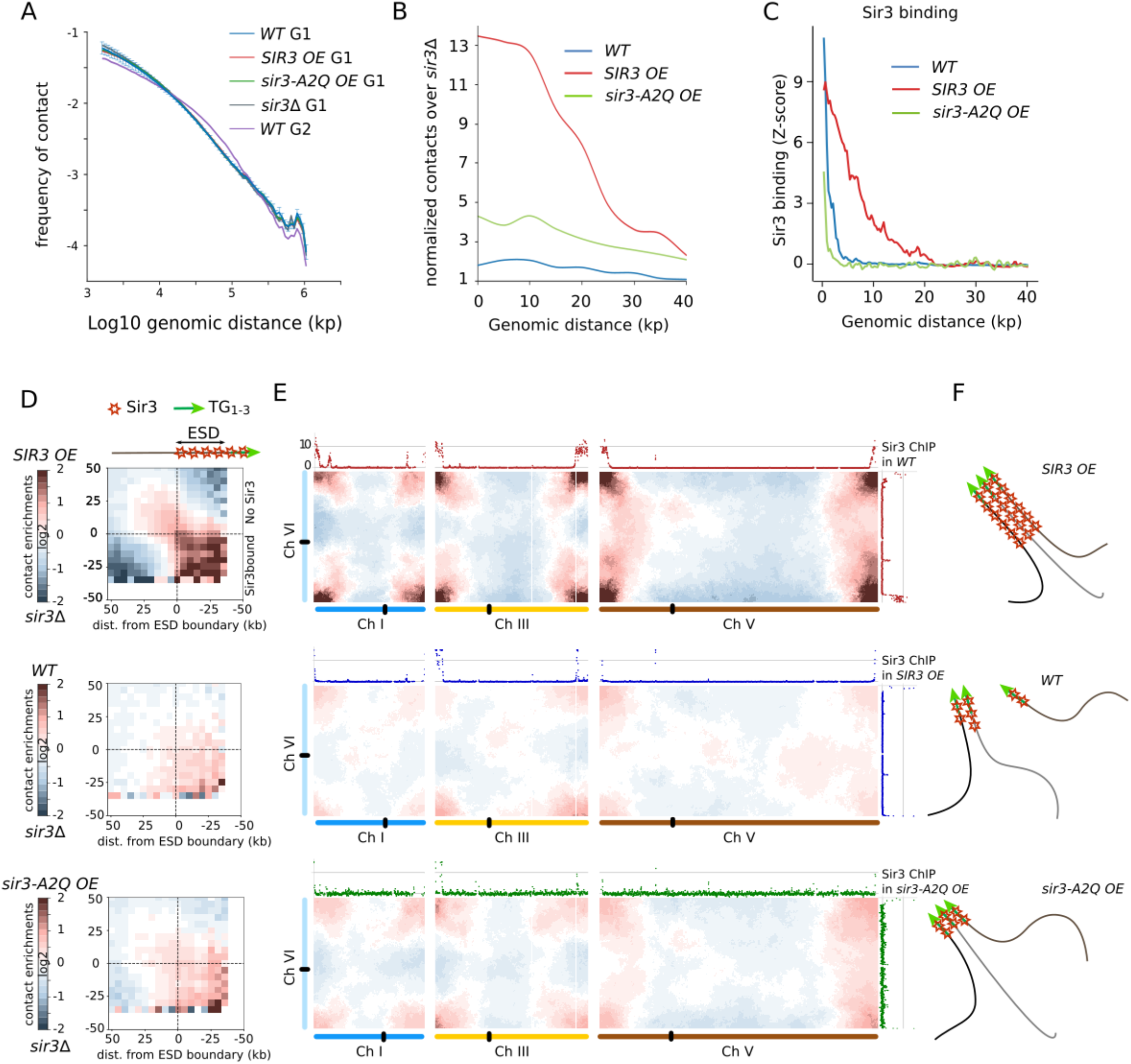
Sir3 spreading coincides with telomere-telomere contacts. (**A**) Left panel: contact probability as a function of genomic distance Pc(s) (log scale) for different strains either in G1 or G2. Right panel: cumulated normalized contacts over 40kb subtelomeric windows (0 = X-core elements). (**B**) Pile-up contact maps ratio for various pairs of strains of subtelomeric windows centered on the extended silent domains (ESD; dotted line) determined in the *SIR3 OE* condition. The windows extend over 50 kb in both direction from the ESD. (**C**) Sir3 enrichment over 40 kb subtelomeric windows. (**D**) Representative interchromosomal ratio maps of the same pairs of strains as in (**C**) processed through Serpentine binning. Sir3 enrichment observed by ChIP in the indicated onditions is plotted along the top and right axis. Right: schemes illustrating the behavior of telomeres and subtelomeric regions in the *SIR3 OE*, *WT* and *sir3-A2Q OE* conditions.

We then focused on telomere-telomere contacts. We plotted the ratio of inter-subtelomere contacts between a given strain and a *sir3*Δ strain as a function of the distance to the telomeres (Figure 2B). The ratio with the signal in the *sir3*Δ strain measures the effects of the Sir3 protein on the contact probability and automatically conceal the Sir3-independent biases present in all strains, due to the high variability of the alignment coverage in subtelomeric regions. For both the wild-type or the strain overexpressing Sir3-A2Q, the ratio was the highest near the telomere and decreased slowly with the distance from the telomere. In good agreement with our microscopy data, inter-subtelomeres contacts were higher in the Sir3-A2Q overexpressing strain than in the wild-type. Sir3 overexpression led to an even higher ratio of inter-subtelomere contacts that decay slowly with the distance to telomere over the first 10kb before decreasing sharply up to 35 kb towards the centromeres. The extent of high level of inter-subtelomere contacts is comparable to the extent of Sir3 spreading as analyzed by ChIP (Figure 2C), suggesting that Sir3 bound regions mediate these contacts.

Indeed, in wild-type cells, Sir3 is detected bound to DNA 2.6 kb away on average from the last telomeric element. In strains overexpressing Sir3, it can be detected as far as 20 kb away from the telomere, generating large Sir3-bound regions, dubbed extended silent domains (ESD, (Hocher et al., 2018) and Figure 2C). In contrast, Sir3-A2Q detection is limited to the TG repeats, even when overexpressed.

To assess the contribution of Sir3 binding to trans-subtelomere contacts, subtelomeric contact maps of strains either overexpressing *SIR3* or *sir3*Δ were centered on the ESD position and piled-up. We then computed the mean matrix, calculated over all these submatrices, of the differential contacts in *SIR3* overexpressing versus *sir3*Δ strains (Figure 2D top panel). This matrix conveys the cumulated amount of inter-chromosomal contacts mediated by Sir3 between subtelomeric loci, as a function of their distance from the respective ESD boundaries. We observed a general increase of trans-subtelomere interactions for all loci located between the telomeres and the ESD boundaries, which we identify as the signature of the telomere hypercluster. Passed the ESD border, trans-interactions decrease sharply, further suggesting that in *SIR3* overexpressed conditions, ESD boundaries are placed at the surface between two different nuclear compartments well defined by their Sir3 occupancy. We also notice a persistence of interactions along the diagonal reflecting weaker interactions between regions adjacent to the ESDs that are possibly maintained in close proximity by the constraint imposed by the telomere hypercluster to the chromatin polymeric backbone. In wild type cells or Sir3-A2Q overexpressing cells we observed a progressive decrease of interactions from the telomere proximal region to the telomere distal region, probably reflecting the bonding of subtelomeres through shorter regions upstream of the ESD boundary (Figure 2D and 2S). Also, in WT cells, the intensity of the interaction is generally lower, reflecting the presence of more than one telomere cluster, possibly due to heterogeneity at the single cell level as a result of random encounters of different couples of telomeres (Figure 1A).

At the individual chromosomes level, to visualize the correlation between Sir3 spreading and contacts between different subtelomeres, we built differential contact maps at an higher level of details using the serpentine algorithm (Baudry et al. 2020), which adapts the resolution on each position of the contact map to the data coverage (Figure 2E, see Figure S2 for maps comprising all chromosomes). The serpentine map comparing the *SIR3* overexpressing cells with the *sir3*Δ showed a strong increase of inter-subtelomeres contacts between ESDs of chromosome VI and chromosomes I, III and V (dark red regions at the corners of the maps). As a consequence, those subtelomeric regions make less contact with non-subtelomeric regions. In the wild-type vs *sir3*Δ serpentine maps we observed a similar, though less pronounced, trend. Thus, contacts between subtelomeric regions coincide with Sir3 binding, suggesting that Sir3 binding is the epigenetic information setting, on the linear chromosomes, the boundary of the telomere clusters in 3D.

### Sir3 but not Sir3-A2Q accumulates at the rDNA upon overexpression

We next asked whether non-subtelomeric Sir3 bound regions interact with subtelomeres. Notably, Sir3 binds to the rDNA through a mechanism that requires Sir2 deacetylase activity (Hoppe et al., 2002; Radman-Livaja et al., 2011). In agreement with this work, an endogenously expressed GFP-tagged version of Sir3, colocalized with an endogenously expressed mCherry-tagged rDNA bound protein Net1, as shown by live microscopy in Figure 3A.

**Figure 3:**
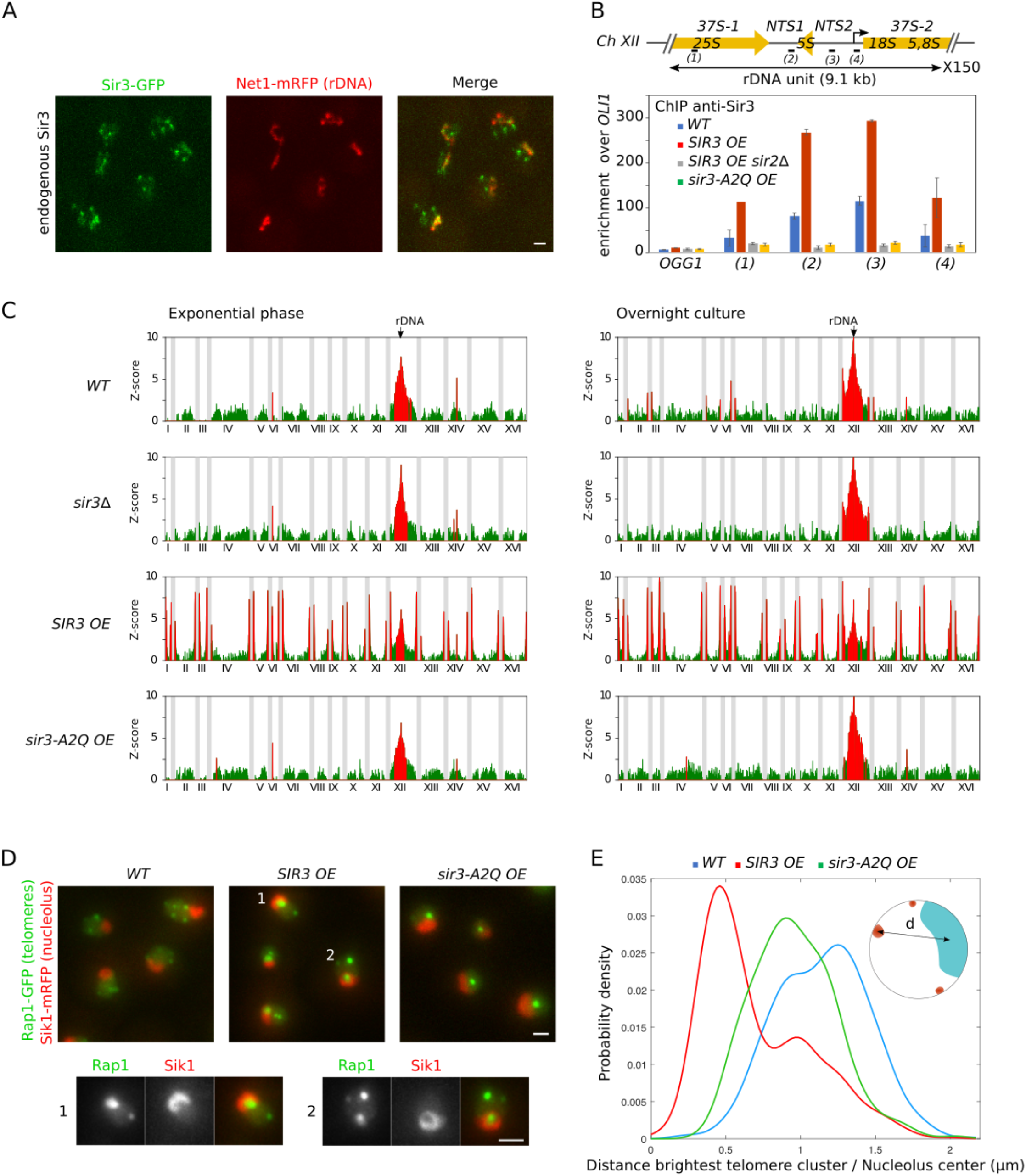
Sir3 but not the silencing dead mutant, Sir3-A2Q, associates with the rDNA and promotes rDNA-telomere contacts. **(A)** Representative fluorescent images of double tagged Sir3-GFP / Net1-mRFP strains expressing either endogenous (yAT2803) or high levels of Sir3 (yAT3113). Cells were grown in synthetic complete medium with 2% glucose and imaged in exponential phase. **(B)** The *S. cerevisiae* rDNA locus is composed of ~150 tandem copies of a 9.1 kb repeating unit, each encoding two transcribed region RDN5 and RDNA37 (comprising RDN18, RDN5.8 and RDNA25 genes). The graph represents Sir3 occupancy along the rDNA locus probed by ChIP-qPCR using an anti-Sir3 antibody (Ruault et al., 2011) in WT (yAT232), *GAL1p-SIR3* (yAT208), *GAL1p-Sir3-A2Q* (yAT1205), and *GAL1p-SIR3 sir2*Δ (yAT772) strains. Primer pair 1 amplifies a region of the RDN25 locus, primer pair 2 amplifies a region in NTS1 (Non-Transcribed Spacer 1) region, primer pair 3 amplifies a region in NTS2 and primer 4 amplifies a region in the ETS1 (External Transcribed Spacer 1) region (see table S2 for primer sequences). Strains were grown in YPGal for 48 hours. The bar graph represents the Sir3 enrichment over the mitochondrial locus *OLI1*. Error bars show s.e.m of three independent experiments, each analyzed in triplicate qPCRs. **(C)** 4C extraction profiles representing contact maps between the rDNA repeats and the rest of the genome, profiles are shown for all mutants in both exponential and overnight cultures. Red highlighted loci correspond to bins with z-value greater than 2.5. **(D)** Representative fluorescent images of a double tagged strain Rap1-GFP / Sik1-mRFP in strains expressing endogenous level of Sir3 (yAT340), high levels of Sir3 (yAT341) or high levels of the separation of function mutant Sir3-A2Q (yAT1198) after an overnight culture in complete synthetic medium with 2% galactose. Magnification of representative nuclei (1 and 2) are presented at the bottom of the panel. **(E)** Distance between the brightest Rap1-GFP cluster and the nucleolus center is plotted for a wild-type (yAT340, n=581), a strain overexpressing Sir3 (yAT341, n=627) and a strain overexpressing Sir3-A2Q (yAT1198, n=590) using the Nucloc software (Berger et al., 2008). Cells were grown in complete synthetic medium 2% galactose overnight before imaging. Scale bar is 1 μm in all panels.

To identify precisely the region bound by Sir3 in a rDNA repeat unit, we performed ChIP experiments (Methods). Sir3 was found preferentially enriched in the non-transcribed regions (*NTS1* and *NTS2*) and, to a lesser extent, around the *RDN25* transcriptional unit and the promoter of the 37s rDNA precursor transcript (Figure 3B). Probing Sir3 binding in a strain overexpressing *SIR3* we observed a strong accumulation of Sir3 along the rDNA unit, with a stronger signal at *NTS1* and *NTS2*, showing that Sir3 concentration was limiting, binding not only at telomeres but also at the *rDNA* locus. Consistently with previous work (Hoppe et al, 2002) we did not observe Sir3 binding in the absence of Sir2 even upon Sir3 overexpression. In contrast to wild-type Sir3, the Sir3-A2Q mutant that has lost its affinity to bind nucleosomes was not detected at the *rDNA* by ChIP. Together these results suggest that Sir3 accumulates at the rDNA through its affinity for unacetylated nucleosomes resulting from Sir2 activity (Figure 3B).

### Sir3 promotes interaction of the telomeres with the rDNA locus

A 4C like analysis using the rDNA locus as a viewpoint (Methods) showed that when Sir3 is expressed at its endogenous level in exponential phase, the rDNA locus makes significant contacts with only two loci outside chromosome XII. When the rDNA locus was less active (overnight cultures), we observed that contacts between the rDNA and other regions of the genome increased to 11, corresponding mostly to rDNA-telomere interactions (10 out of 11). In contrast to the interactions observed in exponential phase these interactions were lost in the absence of Sir3 (Figure 3C). These observations suggested that the activity of the rDNA prevents those Sir3 dependent contacts during exponential phase. In strains overexpressing Sir3, the *rDNA* locus showed statistically significant interactions with all telomeres during exponential phase, showing that the increased amount of Sir3 at both types of loci favored their trans-contacts.

When Sir3-A2Q was overexpressed, the loci in contacts with the rDNA remained similar to WT exponentially growing cells, but significant contacts with telomeres were not detected after an overnight culture. Therefore, rDNA-telomere contacts required Sir3 binding at the rDNA. Note that these analyses are averaged over a cell population, and don’t necessary reflect the contacts of the rDNA in each independent cell of the population.

To observe those contacts at the single cell level, we acquired images of cells expressing Rap1-GFP (telomere foci) and Sik1-mRFP (nucleolus) in strains expressing different *sir3* alleles. Images were taken from overnight cultures cells enriched in rDNA-telomere interactions. In a wild-type strain, telomere foci are located at the nuclear periphery and don’t interact with the nucleolus (Figure 3D). On the opposite, in a strain overexpressing Sir3, the telomeres group together and position in close contact with the nucleolus. More specifically, the telomere hypercluster is set within the nucleolus, excluding the Sik1 protein (magnification on the bottom of Figure 3D). The Sir3-A2Q mediated telomere hypercluster, on the other hand, didn’t interact with the nucleolus, in agreement with Hi-C maps. We next measured the 3D-distance between the brightest telomere clusters and the nucleolus at a single cell level using NucLoc (Berger et al. 2008). The distance between the brightest cluster of telomeres and the center of mass of the nucleolus was shorter in strains overexpressing Sir3 (0.45 μm) compared to wild type strain (1.2 μm) (Figure 3E). In strains overexpressing the Sir3-A2Q mutant, the distance was intermediate (0,9 μm) reflecting the localization of the hypercluster at the center of the nucleus without interacting with the nucleolus, again in agreement with our 4C maps.

Also in agreement with our observations with the 4C plots, the physiological state of the cells influence the level of association between the telomere hypercluster and the nucleolus (Figure S3A). Indeed, the two subnuclear compartments that are tightly associated after an overnight culture in *SIR3* overexpressing cells dissociate upon dilution in fresh medium and would reassociate when cells reach saturation (Figure S3B).

Another factor that influenced the telomere hypercluster-rDNA contact was the number of rDNA repeats: as we observed a much less frequent interaction in cells with a short rDNA (25 copies) compared with cell with high copy number (190 repeats, Figure S3C) in the Sir3 overexpressing mutant.

From this we conclude that the amount of Sir3 bound at the rDNA favors its interaction with telomeres. However, a high transcriptional activity at the rDNA prevents these interactions.

### Sir3 overexpression compacts the rDNA and inverts the spatial organization of the nucleolus

Since Sir3 has a strong impact on the spatial organization of the telomeres, we asked whether the accumulation of Sir3 at the rDNA locus could modify its spatial organization. In wild-type cells the rDNA labelled by Net1-GFP appears as a filament (Figure 4A-B) as previously reported (Straight et al. 1999). In Sir3 overexpressing cells, rDNA compaction varies with the physiological state of the nucleolus (Figure S4A), with an increasing compaction from early to late exponential phase. As expected from cells grown overnight, the compacted rDNA is overlapping with the telomere hypercluster in most cells (Figure 4A bottom panel). In contrast, cells overexpressing the Sir3-A2Q mutant that cannot bind the rDNA showed a filamentous rDNA (Figure 4A) suggesting a direct role for Sir3 in compacting the rDNA.

**Figure 4:**
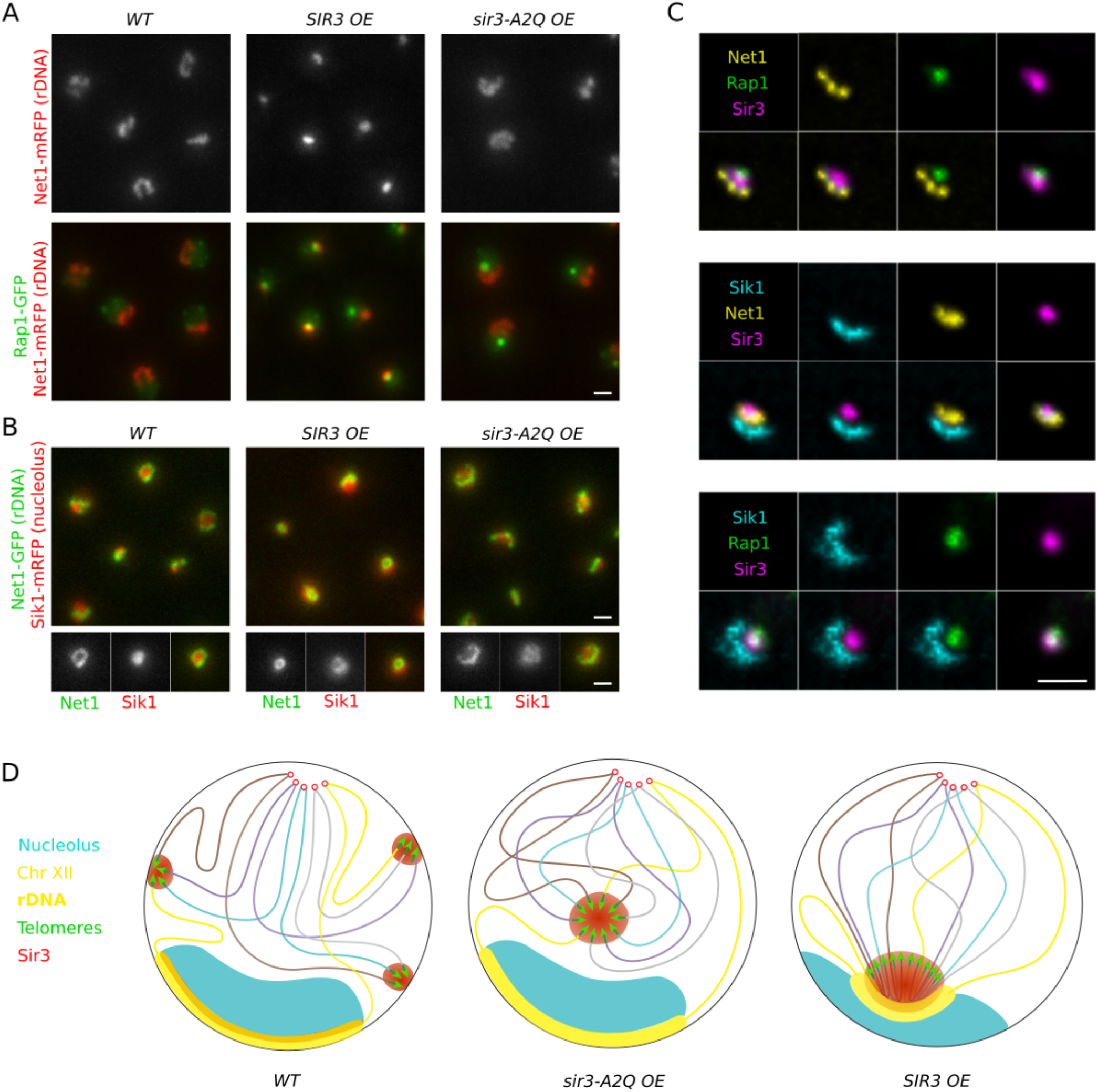
Sir3 overexpression impacts on the rDNA spatial organization and its compaction. **(A)** Representative fluorescent images of Rap1-GFP / Sik1-mRFP strains expressing either endogenous level of Sir3 (yAT3729), high levels of Sir3 (yAT3730) or high levels of the Sir3-A2Q mutant (yAT3733). Cells were grown in synthetic complete medium with 2% galactose and imaged after an overnight culture. **(B)** Representative fluorescent images of a double tagged strain Net1-GFP / Sik1-mRFP in strains expressing endogenous level of Sir3 (yAT1004), high levels of Sir3 (yAT1008) or high levels of the Sir3-A2Q mutant (yAT1541). Cells were grown in synthetic complete medium with 2% galactose and imaged after an overnight culture. In the bottom panels, close-up of selected nuclei from the field are shown (GFP-channel, RFP channel and merge). **(C)** Representative fluorescent deconvolved images of triple tagged strains, individual and merge channels are shown. Cells were grown overnight in synthetic complete medium 2% glucose and are all expressing high Sir3 levels (Sir3 under the control of the constitutive GPD promoter). From the Top to the Bottom: Rap1-GFP Sir3-mCherry Net1-BFP2 (yAT3901), Net1-GFP sir3-mCherry Sik1-BFP2 (yAT2213) and Rap1-GFP Sir3-mCherry Sik1-BFP2 (yAT3666). **(D)** Schema representing the nuclear organization of a wild-type, a strain overexpressing the Sir3-A2Q separation of function mutant and a strain overexpressing the wild-type Sir3. Scale bar is 1 μm in all panels.

We next investigated the consequence of overexpressing Sir3 on the organization of the nucleolus by imaging strains expressing the rDNA bound protein Net1 labeled with GFP, as well as Sik1-mRFP, involved in pre-rRNA maturation and thus staining the nucleolus. The rDNA filament was surrounding the Sik1 signal in wild-type cells or cells overexpressing Sir3-A2Q (Figure 4B). This organization was inverted in Sir3 overexpressing cells, with the Net1-GFP signal located in the center of the nucleolus and surrounded by the Sik1 signal. Both the compaction of the rDNA and the inversion of the nuclear organization were independent of the formation of telomere hyperclusters as they were observed in *sir4*Δ cells overexpressing Sir3 (Figure S4B-C).

Finally, to get a better idea of the interaction between the telomere hypercluster and the nucleolus when they interact, *i.e.* when the activity of the nucleolus is low (overnight cultures), we built different combination of triple labelled strains that allowed us to image the telomere hypercluster (Rap1), Sir3, the rDNA (Net1) and the nucleolus (Sik1) in Sir3 overexpressing strains (Figure 4C). In those conditions, the Sir3 signal from the telomeres was indistinguishable from the rDNA bound one, corresponding to a large focus that overlapped on one side with the Net1 signal, and on the other with the Rap1 signal (figure 4C upper panel and S4D) with no overlap between Rap1 and Net1 signals. Furthermore, the Sir3 signal overlapping with Net1 was always facing the Sik1 signal (i.e. the nucleolus) (Figure 4C, middle panel and Figure S4D), while the Sir3 signal colocalizing with Rap1 was found at the opposite of the nucleolus (Figure 4C lower panel and S4D). Therefore, the subtelomeres (enriched in Sir3 and deprived of Rap1) are in contacts with the Sir3-bound rDNA, while the extremity of the telomeres (enriched with Rap1) protrudes towards the nuclear interior.

Those results led us to propose the following model presented in Figure 4D: in strains overexpressing Sir3, Sir3 binds and coalesces the telomeres into a hypercluster, while accumulating at the rDNA as well and increasing its compaction. When the activity of the nucleolus is low, the regions of the rDNA that are Sir3-bound interact with the telomere hypercluster bringing the rDNA and the telomeres together in a “super” Sir-enriched subcompartment. The part of the nucleolus dedicated to pre-RNA processing is excluded from this subcompartment and is self-organizing between the Sir3 enriched subcompartment and the nuclear periphery.

### Ectopic Sir3-bound regions coincide with long range and trans-contacts

In addition to the telomeres and the rDNA, small peaks of Sir3 binding are also detected by ChIP at a few discrete sites located internally on chromosome arms (Hocher et al. 2018; Mitsumori et al. 2016; Sperling and Grunstein 2009; Takahashi et al. 2011). Although some of these sites were shown to be ChIP artefacts (Teytelman et al. 2013), others were not, because no enrichment was seen in the Sir3-A2Q expressing strain and Sir3 enrichment increased upon Sir3 overexpression (Hocher et al. 2018).

The serpentine maps revealed that some of those internal sites are also in contact with other Sir3-bound loci, especially in cells that have slowed down cell division (*i.e.* overnight cultures). This was clearly visible on the serpentine map of chromosome I and VI when comparing cells overexpressing Sir3 to the ones overexpressing the Sir3-A2Q mutant protein, that is not recruited at these internal sites. Especially, the *YAT1* and *IGD1* genes, respectively located on chromosome I and VI (38 kb and 60 kb from the closest telomere), showed increased contact frequencies with the subtelomeres belonging to the same chromosome (Figure 5A). This was also true for the *YDL007C-A* gene located 13 kb from the centromere of chromosome IV, which interacts with the subtelomere 4L thus counteracting the global Rabl-like organization of this chromosome arm. Indeed, the serpentine map also revealed increased cis-interactions along the left arm of chromosome IV, possibly reflecting the loop formed between *YDL007C-A* and the telomere. Further, we observe increased contact frequencies between the *SIR3* gene, where Sir3 is recruited upon overexpression, and the subtelomeres located 56kb downstream on the right arm of chromosome XII (Figure S5B), possibly hinting on a negative feedback mechanism regulating *SIR3* expression. However, not all Sir3 bound site showed preferential interactions with telomeres, maybe owing to other constraints associated with cis-associated loci. Some of the internal Sir3 bound sites that interact with the subtelomeres of their own chromosome in *cis* also interact in trans with other subtelomeres. Figure 5B is presenting the contacts between chromosomes IV, V and VI and the rest of the genome and is emphasizing the ability of internal Sir3 bound sites to contact any subtelomeres of the genome. No other trans-contacts outside Sir3 bound regions was observed in these maps. However, Hi-C maps of the *sir3*Δ strain shows apparent contacts between the *FLO1, FLO9* and *FLO11* genes located respectively in telomere-proximal regions of chromosomes I right arm, I left arm, and IX right arm (Figure S5A). Further analysis at the read level revealed that the apparent interaction between *FLO1* and *FLO9* could owe from an alignment artefact, while this is not the case for *FLO1-FLO11* interaction (Figure S5B). In strain expressing Sir3 or Sir3A2Q, this interaction is probably also occurring but almost impossible to distinguish from Sir3 mediated interactions between subtelomeres.

**Figure 5:**
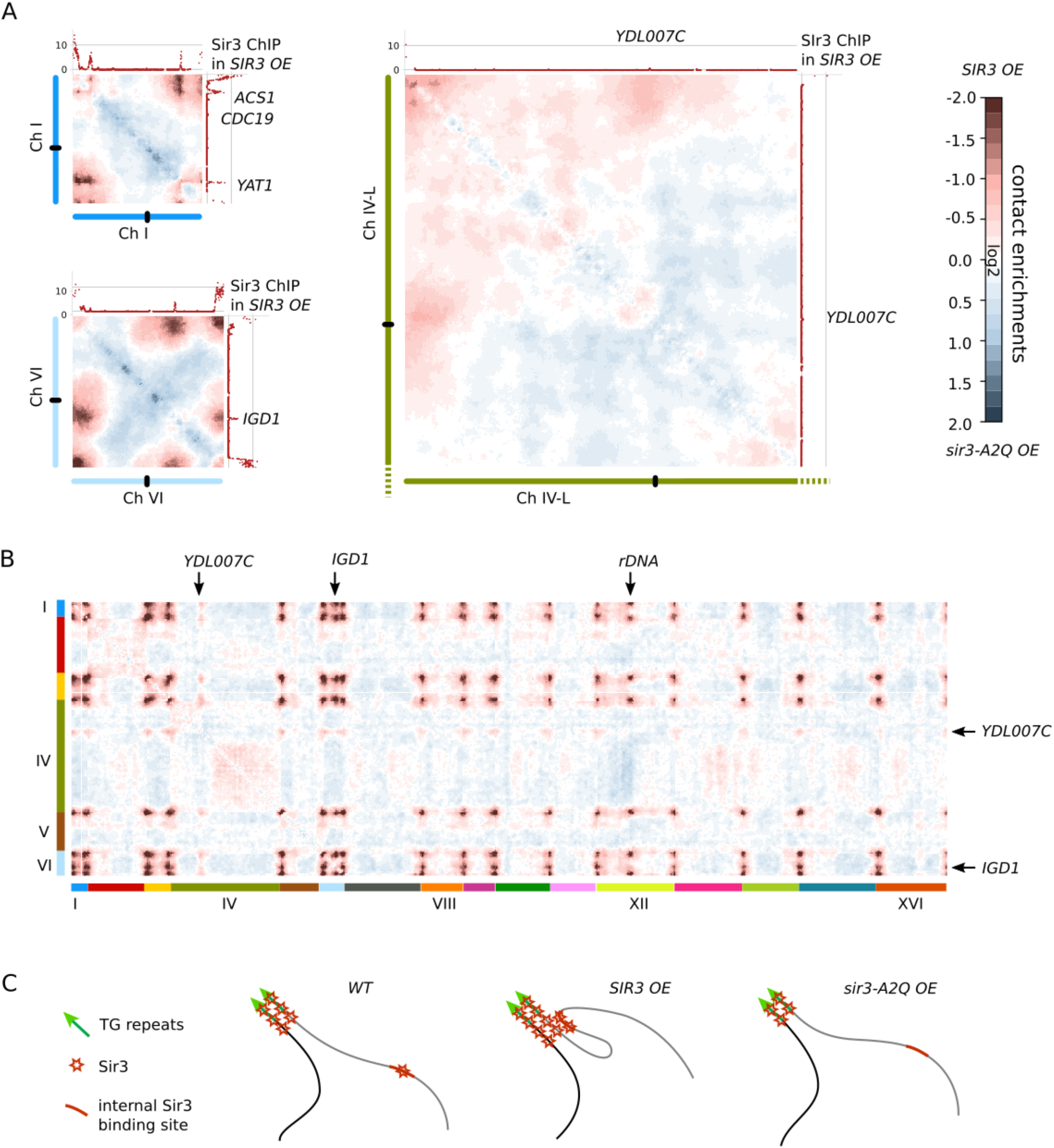
Sir3 bound regions coincide with long range contacts. (**A**) Intrachromosomal ratio plots of chromosome I, VI and IV-L contact maps generated in *sir3a2q OE* and *SIR3 OE* conditions, and processed by Serpentine binning. The ChIP deposition profile of Sir3 in OE conditions are plotted along the top and right axis. Intrachromosomal loci enriched in Sir3 are indicated by the closest gene name. (**B**) Serpentine binning plots recapitulating the enrichment in contacts in *SIR3 OE* conditions respect to *sir3-A2Q O*E conditions, showing chromosomes I,IV,V and VI interacting with the rest of the genome. (**C**) Schematic representation of the behavior of internal Sir3 binding sites in different conditions.

In summary, we observed that upon Sir3 overexpression, increased Sir3 binding at internal sites smaller than 4kb is accompanied by increased interactions between these sites and subtelomeric regions suggesting that Sir3 is directly involved in these chromosome contacts. We do not observe increased interactions between these sites and subtelomeres when comparing the wild type and *sir3*Δ strains. However, we detect an increased compaction of those loci resulting in a higher frequency of contact between those sites and adjacent regions (Figure S5C). It is possible that Sir3 retains a role of segregation and silencing on those sites without affecting the large-scale chromosome architecture in the tested wild type conditions. Alternatively, increased bridging between these loci and telomeres might be present in the cell population, but not enough significant to be detectable by our assay.

### Artificial arrays of Sir3 are sufficient to promote trans-interactions

To directly test whether Sir3 is sufficient to bond 2 ectopic loci together, independently of silencing and clustering of telomeres, we hijacked the GFP-LacI / LacO array system to target Sir3 to specific loci harboring LacO arrays by expressing GFP-LacI fused to Sir3. The fusion of GFP-LacI in N-terminus of Sir3 is abolishing its silencing function, but its clustering function remains fully functional (Figure 6SA-B). LacO arrays were introduced at *LYS2* (chr. II) and *LEU2* (chr. III) loci in a strain unable to form telomere foci (*rap1-17 sir3*Δ) to prevent the recruitment of GFP-LacI-Sir3 to telomeres. We measured the pairing of the two arrays in different strains after an overnight culture. In a strain expressing the GFP-LacI, the two foci were paired in 6% of the cells. When we expressed GFP-LacI-Sir3 the percentage of paired loci increased to 28%. This increased pairing was independent of Sir2 activity as 25% of the cells had a paired array when grown in the presence of the splitomicin Sir2 inhibitor. The Sir3 induced increased pairing was also independent of Sir4, since the percentage of pairing was similar in a WT and *sir4*Δ strain expressing GFP-LacI-Sir3. Because these experiments were performed in strains expressing a *rap1* allele truncated for the Rap1-Sir3 interaction domain, we can infer that the Sir3 mediated pairing is also Rap1 independent. Therefore, this synthetic approach demonstrated that Sir3 can bond together loci belonging to different chromosomes, independently of its interaction with Rap1, and Sir4, and Sir2 activity, or chromosome context. This strongly suggests that Sir3-Sir3 interactions provide the physical link that holds together Sir3 bound loci.

**Figure 6:**
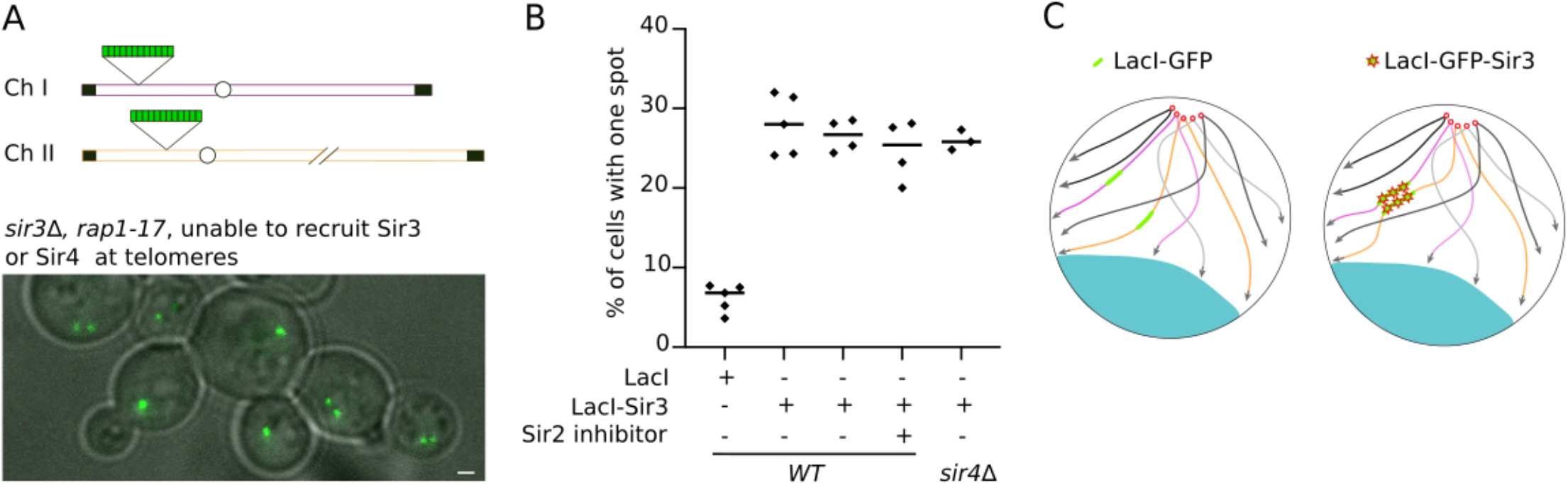
Arrays of Sir3 are sufficient to promote trans-interactions. **(A)** Schema describing the strains used in the assay. *sir3*Δ *rap1-17* strains carrying a LacOp array at the *LYS2* locus on chromosome II and a LacOp array at the *LEU2* locus on chromosome III are expressing either GFP-LacI or GFP-LacI-Sir3. Merge image of a transmitted-light image and the GFP channel fluorescent image of a strain expressing the GFP-LacI-Sir3 construct (yAT1470). **(B)** Graphic showing the percentage of cells with 1 spot in different strains and conditions: GFP-LacI (yAT1476), GFP-LacI-Sir3 (yAT1470) in the presence or in the absence of splitomycin (a Sir2 inhibitor) and GFP-LacI-Sir3 *sir4*Δ (yAT1864). Cells were grown in synthetic complete medium supplemented with 2% glucose to mid exponential phase of growth before being imaged. **(C)** Schema representing the nuclear organization of strains expressing either the GFP-LacI (left) or the GFP-LacI-Sir3 (right).

## Discussion

Here we investigated the impact of the silencing factor Sir3 on the regulation of the yeast genome 3D folding. We show that Sir3 is both necessary and limiting for telomere clustering.

### Sir3 mediates interactions between telomeres

Our Hi-C data clearly show that interactions between subtelomeric regions decrease in the absence of Sir3 and increase upon Sir3 overexpression. These interactions do not require Sir3 spreading since overexpressing Sir3-A2Q, whose binding is limited to the telomeric TG repeats, is sufficient to increase subtelomeric contacts. However, contacts drop rapidly away from telomeres in the Sir3-A2Q strain, whereas in the strain overexpressing the WT version of Sir3 they remain strong over up to 20kb along the subtelomeres. In addition, when Sir3 is overexpressed the increased contacts coincide with Sir3 enrichment along the subtelomeres. This suggests that the subtelomeric contacts in the Sir3-A2Q strain is an indirect consequence of the Sir3-dependent bonding of chromosomes by their telomeres, whereas the spreading of WT Sir3 directly promotes the subtelomeric contacts. Extended Sir3 spreading is also associated with increased interaction in cis possibly reflecting the compaction resulting from heterochromatin formation.

### Sir3 impact on nucleolar organization

Another striking consequence of overexpressing Sir3 is its impact on the organization of the nucleolus. Although Sir3 association with the rDNA has been documented before (Hoppe et al., 2002; Radman-Livaja et al., 2011) its function at this locus remained elusive. Our experiments show that increasing Sir3 association with the rDNA increases its compaction, especially when the rDNA activity is low (i.e. in slowly growing cells). This compaction is accompanied by an inversion of the nucleolar organization: while the rDNA is found at the periphery of the nucleolus in wild-type cells, it becomes nested within the nucleolus upon Sir3 overexpression (Figure 4). In addition, both Hi-C and microscopy show that the rDNA associates with telomeres in a Sir3 dependent manner, in conditions where Sir3 was overexpressed. Although we could not observe this association by microscopy in wild-type cells, such contacts were nevertheless detected by Hi-C for 10 telomeres out of 32 in wild-type cells with low rDNA activity. These observations show that Sir3 influences the overall genome organization not only at the level of subtelomeric regions, but also more broadly.

It is noteworthy that Sir3 recruitment at the rDNA was first reported as a hallmark of aged cells (Kennedy et al. 1997; Sinclair 1997), possibly owing to a lack of sensitivity to detect Sir3 in the nucleolus of young cells. Sir3 was especially visible in the fragmented nucleoli resulting from the instability of the rDNA in old cells. It will be important to test whether this is related to the ability of Sir3 to compact the rDNA and whether this new property of Sir3 impacts on the stability of this locus.

### Tethering Sir3 is sufficient to bridge chromatin loci

In addition to the trans-contacts between large Sir3 covered regions such as subtelomeres and the rDNA, we also observed contacts between Sir3 bound subtelomeric domains and at least 5 discrete, few kb long regions, enriched in Sir3 and positioned internally on chromosome arms (Hocher et al. 2018; Mitsumori et al. 2016; Sperling and Grunstein 2009; Takahashi et al. 2011). Although weaker than contacts between subtelomeres, they can counteract the canonical Rabl-like configuration (where telomeres and centromeres are in opposite sides of the nucleus), drawing together the centromere proximal locus of chromosome 4 and its subtelomeres. These internal sites also interact with subtelomeres from other chromosomes in trans (Figure 5). Although these sites do not show specific contacts with telomeres in wt cells, we could detect an increased compaction of those loci.

While the functional significance of these internal Sir3 binding sites remains to be elucidated, they correspond to intergenic regions or genes that are not expressed in our culture conditions. Of note, one of the most striking examples is the *YAT1* gene that has the highest G+C content of the budding yeast genome (58%) (Chavez et al. 2001) and one of the rare ORF containing a cluster or meiotic double strand break hot spots (Pan et al. 2011). Furthermore, this gene is prone to form R-loops when expressed under the control of the strong *GAL1* promoter (Bonnet et al. 2017). This could be related to our previous findings that Sir3 is recruited at sites of chromatin stress, favoring their perinuclear anchoring (Dubarry et al. 2011). As for the rDNA, it will be important to test whether Sir3 contributes to the genetic stability of these loci.

### Dynamics of Sir3 mediated Long-range contacts

Telomere clusters are dynamic. They can split and fuse (Hozé et al. 2013; Schober et al. 2008), and dissolve during mitosis to reform in G1 phase. We observed that Sir3 overexpression prevents this dissolution probably by stabilizing telomere – telomere interactions. We noticed that telomere clusters are more visible by microscopy as well as by Hi-C in slowly growing cells possibly due to the time required for telomeres to encounter again after mitosis. This is also true for other interactions between Sir3 bound regions, whether the rDNA, internal discrete or synthetic sites. We propose three non-exclusive explanations for this observation. First, some chromatin movements or constraints linked to genome activity could destabilize or prevent Sir3 mediated contacts. Indeed, rDNA-telomere contacts are limited when the rDNA is transcriptionally active, possibly owing to a stronger anchoring at the nuclear periphery when the rDNA is active. Alternatively, the high density of ongoing transcripts could prevent telomeres to contact the rDNA through steric hindrance. Finally, long-range contacts may be limited by the time required for Sir3 bound loci to meet in the nuclear space.

### Mechanisms of Sir3 mediated trans-interactions

Recent work (Gibson et al. 2019) suggests that properties inherent to chromatin, including nucleosomal spacing and histone acetylation status, could promote phase separation phenomena within the nucleoplasm. Reconstituted chromatin undergoes liquid-liquid phase separation (LLPS) in physiologic salt, forming droplets that can be dissolved by acetylating histone tails. Given the presence of the histone deacetylase Sir2 in subtelomeric regions, telomere clustering could result from the LLPS of non-acetytaled chromatin. However, we showed that cells expressing the *sir3-A2Q* allele can form telomere hyperclusters even in the absence of Sir2. Furthermore, the artificial tethering of Sir3 to two distant euchromatic loci is sufficient to bridge them independently of Sir2 activity. These results indicate that the clustering of Sir3 bound loci does not dependent on their chromatin status but rather is directly mediated by direct Sir3-Sir3 interactions. Consistent with this hypothesis, Sir3 carries very well-characterized dimerization domain on its C-terminal part (Oppikofer et al. 2013). In addition, Sir3 C-terminal domain can interact with a more internal part of Sir3 (King et al. 2006). It is therefore very likely that these specific Sir3–Sir3 interactions build-up molecular bridges between Sir3-bound regions, in agreement with *in vitro* studies (Adkins et al. 2009; Georgel et al. 2001; McBryant et al. 2008). In this case, specific Sir3-Sir3 interactions between arrays of Sir3 molecules associated with subtelomeres, the rDNA or internal regions would rather form a dynamic, gel like, network structure rather that a liquid droplet as proposed for other type of nuclear foci (Sawyer et al. 2019). However further work is needed to decipher the physical nature of these foci. Interestingly, heterochromatin foci in mouse cells were first described as liquid droplets (Strom et al. 2017), but a recent study indicates that these foci rather resemble collapsed polymer globules (Erdel et al. 2020). Whether specific heterochromatin factors are responsible for this collapse is not known, but HP1, the functional homolog of Sir3 does not seem required.

We show that Sir3 plays an important, direct role in the colocalization of multiple chromatin loci in budding yeast. How these long-range interactions are regulated in response to environmental cues, eventually reshaping chromosome folding to a larger extent and playing new roles in genetic regulation for instance, has to be further investigated.

## ACKNOWLEDGEMENTS

The authors thank Pr. Masayasu Nomura for sharing strains, Mickael Garnier for his help on image analysis, Julien Mozziconacci and and the members of the Taddei laboratory for helpful discussions. A.T. team was financially supported by funding from the Labex DEEP (ANR-11-LABEX-0044 DEEP and ANR-10-IDEX-0001-02 PSL), from the ANR DNA-Life (ANR-15-CE12-0007), Fondation pour la Recherche Médicale (DEP20151234398), and CNRS grant 80prime PhONeS, the authors greatly acknowledge the PICT-IBiSA@Pasteur Imaging Facility of the Institut Curie, member of the France Bioimaging National Infrastructure (ANR-10-INBS-04). V.S. is the recipient of a Roux-Cantarini Pasteur fellowship. L.L.S. was supported in part by a fellowship from the Fondation pour la Recherche Médicale. This research was supported by funding to R.K. from the European Research Council under the Horizon 2020 Program (ERC grant agreement 771813).

## Materials and Methods

### Media and Growth conditions

Yeast cells were grown either in rich medium (YPD) or in enriched complete synthetic medium (2x final concentration of CSM, MP Biomedicals) supplemented with 2% glucose or 2% galactose (wt/vol). All the strains were grown at 30°C with shaking at 250 rpm. For galactose induction, cells were grown overnight for pre-culture in YPGal medium (yeast extract, peptone, 2% galactose wt/vol) and diluted the next day in the same medium for the experiment. For Sir2 inhibition, splitomycin was added directly to the culture at a final concentration of 62.5 μM for 4 hours. G2/M synchronization was done as in (Lazar-Stefanita et al. 2017). Synchronization at the G2/M transition was achieved by restarting G1 cells in YPD at 30°C for 1 h, followed by the addition of nocodazole (Calbiochem; 15 μg/ml) and incubation for another 2 h at 30°C.

### Strains

The strains used in this study are listed in Table S1 and Table S3. They are all derivatives of W303 (Thomas & Rothstein, 1989) except the ones used for the Hi-C experiment (BY4741, Euroscarf). Gene deletions, insertions of alternative promoters and gene tagging were performed by PCR-based gene targeting (Longtine et al., 1998; Janke et al., 2004).

### Plasmids

pAT234 is an integrative plasmid expressing GFP-LacI-Sir3 under the *HIS3* promoter. This plasmid is derived from a plasmid expressing GFP-LacI-NLS under the *HIS3* promoter (pAFS135,(Straight et al. 1998)) and was built in two steps. First, the STOP codon of GFP-LacI-NLS TAA was mutated to AGA by direct mutagenesis using primer pair am449 (5’ AAAGAAGAAGAGAAAGGTTGCCAGATCTAGAGCGGCCGCCACCGCGGTGG 3’) /am450 (5’ CCACCGCGGTGGCGGCCGCTCTAGATCTGGCAACCTTTCTCTTCTTCTTT 3’) yielding pAT233. The full length *SIR3* gene was amplified by PCR using primer pair am446 (5’ ATAAGAATGCGGCCGCaATGGCTAAAACATTGAAAG 3’) /am448 (5’ AGTCGAGCTCTCAAATGCAGTCCATATTTTTG 3’). The resulting PCR product was digested NotI/sacI and cloned into pAT233 digested NotI/sacI to generate the plasmid encoding the GFP-LacI-Sir3 protein.

### Chromatin Immunoprecipitation experiments

A total of 20 OD600nm equivalent of cells were fixed in 20 mL with 0.9 % formaldehyde for 15 min at 30°C, quenched with 0.125 M glycine for 5 min and washed twice in cold TBS 1x pH 7.6. Pellets were suspended in 1mL TBS 1X, centrifuged and frozen in liquid nitrogen for - 80°C storage. All following steps were done at 4°C unless indicated. Pellets were resuspended in 500 μL of lysis buffer (0.01% SDS, 1.1% TritonX-100, 1.2 mM EDTA pH8, 16.7 mM Tris pH8, 167 mM NaCl, 0.5 % BSA, 0.02 g/L tRNA and 2.5 μL of protease inhibitor from Sigma P1860) and mechanically lysed using a Fastprep instrument (MP Biomedicals) with 0.5mm zirconium beads (Biospec Products): intensity 6, 3 cycles of 30 s with 3 min incubation on ice in between cycles. The chromatin was fragmented to a mean size of 500 bp by sonication in the Bioruptor XL (Diagenode) for 14 min at high power with 30s on / 30s off and centrifuged 5 min at 16000 g. 10 μL were kept to be used as Input DNA. Cleared lysate was incubated overnight with 1 μL of polyclonal antibody anti-Sir3 (Agro-bio). 50 μL of magnetic beads protein A (NEB) were added to the mixture and incubated for 4 h at 4°C on a rotating wheel. Magnetic beads were washed sequentially with lysis buffer, twice with RIPA buffer (0.1% SDS, 10mM Tris pH7.6, 1mM EDTA pH8, 0,1% sodium deoxycholate and 1% TritonX-100), twice with RIPA buffer supplemented with 300 mM NaCl, twice in LiCl buffer (250 mM LiCl, 0.5% NP40, 0.5 % sodium deoxycholate), with TE 0.2% TritonX-100 and with TE. Input were diluted 1/10 with elution buffer (50mM Tris, 10mM EDTA pH8, 1% SDS) and beads were resuspended in 100 μL of elution buffer. A reversal cross-linking was performed by heating samples overnight at 65°C. Proteins were digested with proteinase K (0.4 mg/ml) in the presence of glycogen and the remaining DNA was purified on QIAquick PCR purification columns. Finally, samples were treated with 29 μg/mL RNAse A for 30 min at 37°C and used for quantitative PCR.

### ChIP quantification by quantitative PCR

Quantitative PCR was performed on 1/40 of the immunoprecipitated DNA and 1/3200 of the input DNA. Sequences of interest were amplified using the SYBR Green PCR Master Mix (Applied Biosystems) and the primers listed in table S2. PCR reactions were conducted at 95°C for 10 min followed by 40 cycles at 95°C for 15 s and 60°C for 30 s on a real-time quantitative PCR system (7900HT Fast Real-Time PCR; Applied Biosystems). Each real-time PCR reaction was performed in triplicate. The signal from a given region was normalized to the one from the *OLI1* control locus in immunoprecipitated and input DNA samples. Plots represent the mean value obtained for at least three independent experiments; error bars correspond to SEM.

### Microscopy

Set of images from any given figure panels were acquired using identical acquisition parameters. For all fluorescent images, the axial (z) step is 200 nm and images shown are a projection of z-stack images. Images were acquired on two different systems, either a wide-field microscopy system or a spinning disk system, both of them being driven by MetaMorph software (Molecular Devices). Images of panel 3A, 3E, S3A, S3C, S6B were acquired using a wide-field microscopy system based on an inverted microscope (TE2000; Nikon) equipped with a 100x/1.4 NA immersion objective, a charge-coupled device (CCD) camera (Coolsnap HQ2; Photometrics). A xenon arc lamp (Lambda LS; Sutter Instrument Co.) was used to illuminate the samples. A dual-view micro-imager device, described in Guidi et al., 2015, allows the simultaneous measurement of two-color information on the same sensor. Images of panels 1A, 4A, 4C, S4A and S4D were acquired with the same microscope, using a C-mos camera and another illumination system the Spectra X light engine lamp (Lumencor, Inc). This system allows the fast acquisition of dual-color images when used in combination with a double filter. Images of panels 3D, 4B, 6A, S4B, S4C were acquired on a spinning-disk confocal microscope (Revolution XD Confocal System; ANDOR) equipped with a spinning-disk unit (CSU-X1; Yokogawa), a microscope (Ti 2000; Nikon) with a 100x/1.4 NA oil immersion objective, and an EM CCD camera (iXON DU-885; ANDOR). Images of panel 3A were acquired with an upgraded version of the system allowing the simultaneous acquisition of the GFP and RFP channels using a Tu-Cam module (Andor).

### Microscopy data processing

Deconvolution of images acquired with the wide-field microscopy system was made using the Meinel algorithm in Metamorph (8 iterations; sigma = 0.8; frequency 3; MDS Analytical Technologies).

### Hi-C library generation

Hi-C libraries were generated as described in Lazar-Stefanita et al. (2017), using a DpnII four-cutter enzyme. The Hi-C libraries were sequences on an Illumina NextSeq500 apparatus (2 x 75 high-throughput kits). The PE reads were filtered and aligned according to the protocol described in Cournac et al. (2012). Sequencing libraries are publicly available on the SRA database (accession number SRR12108210 to SRR12108219).

### Visualization of contact maps

Sparse matrices were binned at 5 or 50kb pixel size as indicated in figure legends. Normalization, when applied, used the sequential normalization procedure (SCN; Cournac et al., 2012)) and rasterized using a color-scale.

### Ratio maps and serpentine binning

The log2 ratio of pairs of raw contact maps, binned at 50kb,was computed. The resulting matrix was normalized by subtracting the median of its values. The result was rasterized using a colorscale. Serpentine binning was applied for some of the ratio plots. Serpentine is an algorithm developed to overcome the limits of standard binning with respect to read coverage. When using Serpentine the bin size are chosen non-uniformly on the maps as a function of the sequencing coverage (see Baudry et al., 2020 for validation and details on the program). Filtering was applied to remove speckles to avoid artefacts. The Serpentine ratio allows emphasizing contact patterns otherwise drown in the sampling noise of Hi-C data. Serpentine binning was run independently on contact maps of single or couple of chromosomes, binned at 1250bp resolution, with the following parameters: threshold=70 and minthreshold=7, as discussed in Baudry et al., 2020. Resulting ratio maps were then normalized by subtracting the mean value of their pixels and rasterized using a colorscale centered in zero.

### Contact probability as a function of genomic distance

To compute the intra-chromosomal *P*(*s*) plots, pair of reads aligned in intra-chromosomal positions were partitioned by chromosome arms. Reads oriented towards different directions or separated by < 1.5 kb were discarded to discard self-circularizing events. For each chromosome, read pairs were log-binned as a function of their genomic distance *s* (in kb), according to the following formula: *bin* = floor(log(*s*) /log(1.1)). The *P*(*s*) plot is the histogram computed on the sum of read pairs for each bin. This sum is weighted by the bin size 1.1^(1+*bin*)^ (because of the log-binning), as well as the difference between the length of the chromosome and the genomic distance *s*. The difference acts as a proxy for the number of possible events.

### Inter-subtelomere contacts as a function of the distance to telomeres

The contacts are obtained by comparing the maps obtained in exponential phase of the wild type, the Sir3-overexpression mutant and the Sir3-A2Q mutant, to the maps obtained in the Sir3-delta mutant. For each telomere we obtained, in tabular form, the interactions between the bin containing the X-core element with the sub-telomeres of all the *other* chromosomes. Chromosome III was excluded from the analysis. The table is then reordered as a function of the distance between subtelomere location and the X-core element of the corresponding telomere. The values obtained the aforementioned way in the considered condition are then divided by the values obtained the same way in the Sir3-delta mutant. The mean ratio-table, obtained over all X-core elements and subtelomeres, was then computed. Discarding infinite and NaN values. The resulting table represents the Inter-subtelomere contacts as a function of the distance from telomeres for the considered condition.

### Detection of ESD boundaries

Sir3 binds to telomeres and subtelomeres in the wild type and overexpressed strains, for this reason, the z-transformed values are higher than 1 at the extremities of chromosomes.

For each chromosome, the two ESD boundaries were chosen as the coordinate of the lowest/higher bin in the chromosome with z-transformed value less than 1 in ChIP signal of the overexpressed Sir3 mutant.

### Cumulated ratio-matrices at the ESD boundary

The cumulated ratio-matrices were obtained starting from two raw contact maps binned at 5kb, here named the biological condition and the control. For each couple of ESD boundaries belonging to different chromosomes, excluding chromosome 3, for each matrix, a submatrix was obtained such that the pixel accounting for the direct ESD-ESD interaction is placed at the center. Each of those submatrices is then reoriented such that the telomere-telomere boundary is placed toward the lower right corner, all interactions beyond the telomere-telomere boundary (such as interaction belonging to adjacent chromosomes) are discarded by setting the values to NaN. For each of those submatrices, the log2 of the ratio between the biological condition and the control is obtained through the following formula. Ratio-submatrices are then normalized by subtracting to all of them the same value, computed by the mean of all the pixel in all the ratio-submatrices, excluding NaNs and Infinite values. Finally, the cumulated ratio-matrix is the mean of all ratio-submatrices.

### 4C extraction profiles

From Hi-C matrices, is it possible to extract interactions between a single locus on the genome, with the rest of the genome, by slicing the full rows of the matrix corresponding to the bin containing the locus of interest. If the locus span multiple bins, we sliced multiple rows and then reduce the signal to 1 dimension by taking the sum of the values for each column. In the case of the 4C of the rDNA locus, in order to obtain maximum specificity, the raw data was realigned on a genome with the rDNA removed from the sequence of chromosome 12 and placed on an individual chromosome, stripped of any repeated sequences. The 4C data was then obtained by slicing the full rows corresponding to that individual chromosome, and summing the values of each columns, on the matrix obtained using this special genome as a reference. To detect significant interactions, we plot in green color the z-transform of the 4C signal. The procedure is computed on the logarithm in base 10 of the 4C signal. The median *μ* and the median absolute deviation (*MAD*) was computed. The standard deviation was estimated using the following formula: *σ* = *MAD* / 0.675. The z-transformed 4C is obtained by the following formula: *z* = (= — μ)/σ. Interactions are considered significant, and plotted in red color, according to the following condition: z > 2.5.

## Materials and Methods for Supplementary figure 6

### Silencing assays

For telomeric silencing assays, cultures were grown in liquid YPD and plated in five-fold serial dilutions starting at OD_600nm_ =1 (10^7^ cells /ml) onto appropriate plates. 5-fluoroorotic acid (5-FOA; Zymo Research) plates were prepared by adding 5-FOA to a final concentration of 0.1% to supplemented synthetic medium.

## Supplementary figures

**Figure Sup for Figure 1: High resolution (1250bp bins) normalized contact maps**. Collected are all conditions and mutants considered in this manuscript. For each condition, we provide also a raw contact map (alignment counts per bin) in numpy format; an index pointing to each chromosome start and end bins is attached to the data.

**Figure Sup for Figure 2: Genome-wide ratio maps processed by Serpentine.** Corresponding to the magnification displayed in Figure 2D and Figure 5A and 5B.

**Figure Sup for Figure 3: Association of the telomere hypercluster and the rDNA in strains overexpressing Sir3 is regulated by the physiology of the cell and the size of the rDNA array. (A)** Distance between the brightest Rap1-GFP cluster and the nucleolus center is plotted for a wild-type (yAT1782) and a strain overexpressing Sir3 (yAT1827) measured in different conditions: after an overnight culture (n=757 for yAT1782, n=596 for yAT1827), in early (n=652 for yAT1782, n=546 for yAT1827) and late exponential phase (n=739 for yAT1782, n=513 for yAT1827). Cells were grown in complete synthetic medium 2% galactose. Imaged were analyzed using the Nucloc software (Berger et al., 2008). **(B)** Representative fluorescent images of a double tagged strain Rap1-GFP / Sik1-mRFP overexpressing Sir3 under the GPD promoter (yAT1046) in different physiological conditions: after an overnight culture and 3h, 6h and 9h after dilution in fresh medium. Cells were grown in synthetic complete medium with 2% glucose. Scale bar is 1 μm. **(C)** Distance between the brightest Rap1-GFP cluster and the nucleolus center is plotted for a wild-type (yAT340, n=581, same data than in Figure 3E), for a long rDNA strain overexpressing Sir3 (190 repeats, yAT1778, n= 588) and for a short rDNA strain overexpressing Sir3 (25 repeats, yAT1780, n= 367). Cells were grown in complete synthetic medium 2% galactose. Data were acquired after an overnight culture and analyzed using the Nucloc software (Berger et al., 2008).

**Figure Sup for Figure 4: rDNA spatial reorganization upon Sir3 overexpression is independent of the formation of the telomere hypercluster and is regulated by the physiology of the cell. (A)** Representative fluorescent images of a double tagged strain Rap1-GFP / Sik1-mRFP in a wild-type strain (yAT3729), in a strain expressing high Sir3 levels (yAT3730), in a *sir4*Δ (yAT3765) and in a *sir4*Δ expressing high Sir3 levels (yAT3743). Cells were grown in synthetic complete medium with 2% galactose and imaged after an overnight culture. **(B)** Representative fluorescent images of a double tagged strain Net1-GFP / Sik1-mRFP in wild-type strain (yAT1004), in a strain expressing high Sir3 levels (yAT1724), in a *sir4*Δ (yAT1723) or in a *sir4*Δ strain expressing high Sir3 levels (yAT2124). Cells were grown in synthetic complete medium with 2% glucose and imaged after an overnight culture. **(C)** Representative fluorescent images of a double tagged strain Net1-GFP / Sik1-mRFP in wildtype strain (yAT1004) and in a strain expressing high Sir3 levels (yAT1724). Images were taken in different physiological conditions: after an overnight culture and 5h and 8h after dilution in fresh medium. **(D)** Additional representative cells related to Figure 4C. scale bar is 1 μm for all the panels.

**Figure Sup for Figure 5: Contacts between *FLO1, FLO9* and *FLO11* genes; effects of Sir3 binding on *SIR3;* contact between internal binding sites and subtelomeres in wild type conditions. (A)** Normalized contact maps binned at 5kb, showing the interactions between chromosome I with hitself and chromosome I with IX, in *sir3*Δ strains grown in exponential conditions. Green arrows point to the bins connecting *FLO1, FLO9* and *FLO11* genes. **(B)** Zoom of the maps shown in panel A, for two different classes of aligned reads (same direction vs convergent + divergent reads). Contacts that are result of alignment artefacts, often display a bias toward one of the two class of reads with respect to the other. This can be seen in the *FLO1 vs FLO9* interaction (green arrow left panel), and not in the *FLO1 vs FLO11* interactions (green arrow right panel). Finally, arrow plots show contact maps at read level. The green arrow on the arrow plots of the left panel points to the couple of sites that account for the alignment bias between different read classes. This is localized between a couple of regions that present high degree of homology between the *FLO1* and *FLO9* genes. **(C)** Serpentine maps of chromosome I and VI, comparing Hi-C maps obtained in overnight culture, showing contact enrichments between wild type and *sir3*Δ mutant; cells overexpressing Sir3 to wild type, and cells overexpressing Sir3 to *sir3*Δ mutant. Green arrows represent the position along the diagonal of internal Sir3 binding sites.

**Figure Sup for Figure 6: Trans-interaction are independent of Sir3 silencing function. (A)** Growth assay, telomeric silencing assay at *telVIIL::URA3* were carried out with the following strains: wild-type (yAT69), *sir3*Δ (yAT1010), *pHIS3-GFP-Sir3* (yAT1011), *pHIS3-GFP-SIR3 sir3*Δ (yAT1676), *pHIS3-GFP-LacI-Sir3* (yAT821) and *pHIS3-GFP-LacI-Sir3 sir3*Δ (yAT921). To monitor telomeric silencing at *telVIIL::URA3*, strains were grown in YPD overnight and plated in 5-fold serial dilutions starting at OD600nm= 1 (corresponding to 1 x 10^7^ cells/ml) onto appropriate plates: complete CSM without uracil as a growth control plate and onto a 5-FOA plate to assess telomeric silencing. Decreased growth on the 5-FOA plate reflects the disruption of telomeric silencing (expression of the *URA3* gene at telo *VIIL*). **(B)** Representative images (Transmitted-light image and the GFP channel fluorescent image) of a strain expressing the *pHIS3-GFP-LacI-Sir3* construct (yAT788) grown in complete synthetic medium (exponential phase).

